# Shock and Detonation Waves at an Interface and the Collision of Action Potentials

**DOI:** 10.1101/2020.11.03.366310

**Authors:** Shamit Shrivastava

## Abstract

Action potentials in neurons are known to annihilate each other upon collision, while there are cases where they might penetrate each other. Compression waves that travel within the plasma membrane of a neuron have previously been proposed as a thermodynamic basis for the propagation of action potentials. In this context, it was recently shown that two-dimensional compressive shock waves in the model system of lipid monolayers can nearly annihilate each other upon head-on collision when excited close to a phase transition. However, weaker shock waves showed penetration. In general, once the approximation of small perturbation is not valid, compression waves do not interact linearly anymore. While experiments in lipid monolayers demonstrated this principle, a mechanism remained unclear. In this article, we summarise the fundamentals of shock physics as applied to an interface and how it previously explained the observation of threshold and saturation of shockwaves in the lipid monolayer (all – or – none). While the theory has the same fundamental premise as the soliton model, i.e. the conservation laws and thermodynamics, we elaborate on how the two approaches make different predictions with regards to collisions and the detailed structure of the wave-front. As a case study and a new result, we show that previously unexplained annihilation of shock waves in the lipid monolayer is a direct consequence of the nature of state changes, i.e. jump conditions, within these shockwaves, and elaborate on the consequence of these results for the general understanding of the excitation waves in a thermo-fluids framework.

## INTRODUCTION

The propagation of nerve impulses or action potential is conventionally described as an electrical response of an RLC circuit(1) based on the Hodgkin and Huxley model. The components of the circuit may directly correspond to physiological elements such as membranes or proteins. However, it is widely acknowledged that such a model doesn’t explain a variety of observations made during an action potential, for example the mechanical and thermal changes. There are two ways to address these inconsistencies, first is to try refine the Hodgkin and Huxley model through additional equation to couple the mechanical and thermal changes with the electrical ones(2). The second is to come up with an entirely new description based on first principles, i.e. nerve impulse as a thermofluids phenomenon(3–5). The second approach completely changes the mechanistic understanding of action potentials and provides new interpretations for previous experimental observations. Therefore, the debate is whether a refinement of the model will be sufficient to address the limitations and if it is possible to do so without excluding any essential physics, or are such refinement scientifically inconsistent making a reconciliation through incremental refinements impossible? If the answer is latter, it will change the fundamentals of biological communication not only at cellular level but also how it scales due to the entirely different physics.

The concepts behind nerve impulse as an electromechanical wave that propagate in the plasma membranes were first proposed by Konrad Kaufmann in 1980s(5). Kauffman emphasized the importance of the interface as an independent system in thermodynamics, which has its own state diagrams(6). Therefore, the predictions of the theory were not only applicable to action potentials but interfaces in general, which made it possible to test it even in artificial systems, such as lipid monolayers. In particular, it was explicitly predicted that action potentials should *also* be possible in pure phospholipid systems, and we quote, “*Phase diagrams of phospholipid bilayers and monolayers demonstrate the appearance of a threshold for the onset of a transitions. A thermodynamic force K as a function of arbitrary variable n (e.g. surface pressure in dependence on area per lipid molecule) predicts the following responses*:

1. *Resting state in the fluid lipid phase*
2. *Non-propagative subthreshold excitation*
3. *Transitional all – or – none excitation in the lipid state*
4. *Small increase in amplitude above the threshold of stimulation (saturation).”*

Note that while claiming that propagation of these pulses is adiabatic, it was also clearly stated that upon the completion of excitation cycle, the new state is different from the initial resting state due to the effect of dissipation, hence irreversibility was not ruled out rather that the reason for propagation was reversible, as will be discussed in detail below. In this context, the compression waves in a two-dimensional system of lipid-proteins monolayers at the air/water interface have emerged as an important phenomenon, and indeed all four predictions were shown to be correct over past seven years(7–10).

Lipid monolayers at the air/water interface provide a robust platform for investigating interfacial compression waves, also known as Lucassen waves(11–13). The propagation of these waves is characterized by the interfacial or lateral compressibility of the interface, in analogy with sound waves, hence they are also referred to as two-dimensional sound waves. A natural extension of this analogy would predict that shock waves must also exist in such systems, i.e. propagating waves in which the state of the system changes nonlinearly or discontinuously. Therefore, the physics of shockwaves is deeply relevant to obtain a detailed description of Kauffman’s “transitional all – or – none excitation in lipid states”, i.e. a propagating phase transition, which is by definition discontinuous. Indeed, while testing the predictions of Kauffman for lipid monolayers, the experiments provided new observations, such as splitting of these waves(9), that were found to be consistent with the physics of shock and detonation waves(14). These observations provided detailed mechanistic insights into the possible origin of all the nonlinear characteristics of action potentials, such as threshold, saturation, and annihilation upon collision, all of which have now been observed in lipid monolayers.

The suggestion that compression waves or sound waves can be a potential basis for action potentials has not been generally accepted, despite being around for a long time. We see three primary reasons for it: (a) the strongly nonlinear properties of action potentials which are considered highly unusual for an acoustic phenomenon, but are not unusual in the field of nonlinear acoustics, (b) these ideas invoke the principles of acoustic physics in their most general form, which for example means that the acoustic wave in not just a density wave but a propagating adiabatic perturbation in all the thermodynamic variables, (c) the nature of phase transition itself in a multidimensional phase space can become very non-intuitive, i.e. the phase transition may appear different or not at all depending on the thermodynamic process and the choice of variable(15). For example, during quasi-static processes in real membranes, the state has been measured to change nonlinearly as a function of bulk pH (at constant temperature) but not temperature (at constant bulk pH)(16, 17). In contrast, during an acoustic wave, both pH and temperature change (at constant entropy).

However, these ideas have started becoming more intuitive with a clear demonstration of action potential like nonlinear properties by sound waves in artificial model systems such as a lipid monolayer. While the search for nonlinear waves in lipid monolayers was originally motivated by the predictions of Konrad Kaufman, our experiment designs were also deeply influenced by the quantitative predictions of Heimburg and Jackson (18), who derived a wave equation for electromechanical waves in lipid membranes. While fundamentally the lipid monolayer experiments agree with the premise of soliton model, i.e. the importance of conservation laws and thermodynamics, there are significant differences in the details. The experiments in the lipid monolayer instead led us to the classical framework of shock and detonation physics(19–22), where acoustic waves near phase transitions have previously been investigated in extensive theoretical and experimental details(19, 23–26).

Therefore, the objective of this article is to introduce the basics of the general thermo-fluids framework, which includes all kind of waves, including detonation and deflagration waves. So even if Hodgkin and Huxley equations represent the phenomenon correctly, an equivalent description must be possible in the framework. Finally, the collision of all – or – none waves in lipid monolayers is presented as a case study on how to apply this framework to obtain new insights into a previously unknown mechanism.

## BASIC CONCEPTS IN SHOCK PHYSICS

Shock physics deals with the propagation of discontinuities in a medium, which are unavoidable during any perturbations and thus provides a broad framework that became necessary to resolve the difficulties in the theory of sound(27). The reversibility of sound waves is only true for infinitesimal amplitudes, which might appear to be an idealisation but can in fact be observed experimentally. For example, when monochromatic light of a certain wavelength interacts with a material, it results in Brillouin scattering which appears as a new peak in the neighbourhood of the incident wavelength. The shift in the peak is directly related to the speed of sound in the material as it results from momentum transfer between phonons and photons. Thermodynamically, the dielectric *ξ*(*p*, *s*) of a material when expressed as function of pressure *p* and specific entropy *e* explains Rayleigh scattering as the fluctuations in *ξ* due to fluctuations in entropy at constant pressure, and Brillouin scattering due to fluctuations in pressure at constant entropy, i.e. acoustic fluctuations(28). As an aside, the coupling between the Rayleigh and Brillouin scattering, which occurs in the presence of nonlinearity or relaxation, is how action potentials are believed to couple to ion channels in the thermodynamic framework(29).

The infinitesimal amplitude acoustic waves or acoustic fluctuations are reversible and adiabatic, i.e. isentropic, however macroscopic acoustic perturbations come under the category of finite amplitude waves and are not strictly reversible. In 1800s, these waves presented a significant difficulty in the theory of sound as it became clear that conservation laws and the isentropic equation of state cannot be satisfied simultaneously for finite amplitude waves(27). The difficulty was resolved with the realization that finite amplitude waves follow a different state trajectory than an adiabat, which came to be known as Hugoniots(19). The trajectory essentially represents the locus of final states across a shock jump of different amplitudes starting from a given initial state. The situation is elaborated for a 1D shock represented by the jump Δ in the state variables across the wave front as shown in figure 1. The state jumps from initial state (*p*_0_, *v*_0_) with pressure *p*_0_ and specific volume *v*_0_, to the final state (*p*_1_, *v*_1_). The flow velocity changes from *u*_0_ to *u*_1_, and the specific energy changes from *e*_0_ to *e*_1_. Then the conservation laws in the inviscid limit (no losses to the environment and no thermal conduction along the direction of propagation) can be written as (details from (19));

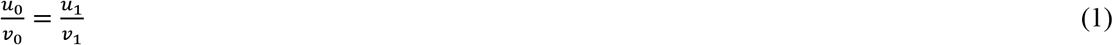

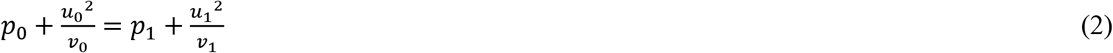

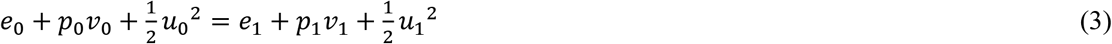

**Figure 1.**
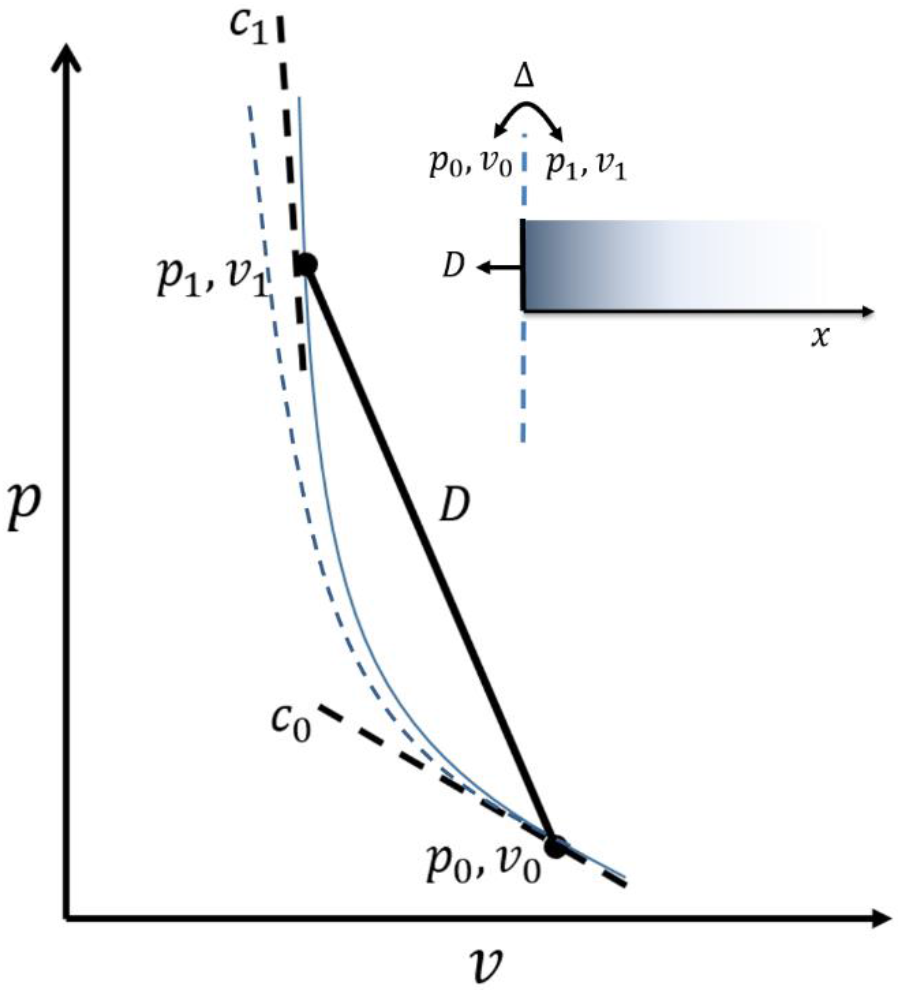
Basics of shock physics: the difference between Hugoniot and adiabatic state diagram. *The state changes across a shock jump propagating in the* – *x direction are shown on the* (*p*, *v*) *diagram of a material. The two states are connected by the straight line of Todes (solid black) which has a slope that is relation to the speed of the wave-front D*. *D* > *c*_0_ *i.e. the speed of sound in the initial state and* < *c*_1_*, i.e. the speed of sound in the final state. The locus of final states thus lies on a state diagram known as Hugoniot (solid blue curve), which diverges from the adiabatic state diagram (dashed blue curve) as the amplitude increases*. *Note that the temperature and pressure are higher in an Hugoniot compared to a adiabatic state change, because the heat from irreversibility stays trapped within the shock wave*.

By combining eq.1 and 2 we get;

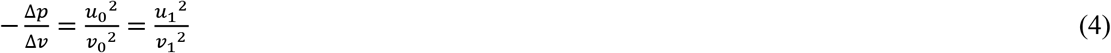

Similarly, by eliminating the velocities *u*_0_ and *u*_1_ from eq. 3, we obtain a relation between the thermodynamic quantities before at after the shock jump, i.e. the Hugoniot equation;

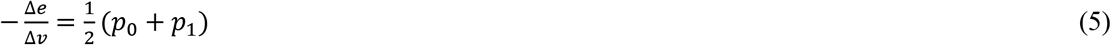

Notice that in the infinitesimal limit, i.e. Δ*v* → 0 we get

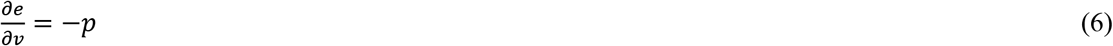

Which implies adiabatic change of state where *e* ≡ *e*(*s*, *v*) and eq.4 then becomes the definition for the speed of sound;

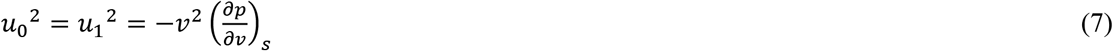

Therefore, the above equations basically represent a generalisation of the theory of infinitesimal sound waves. If the medium is at rest and the observer moves along with the steady wave-front of velocity *D* then eq. 7 can be written in the generalised form for a finite amplitude wave as;

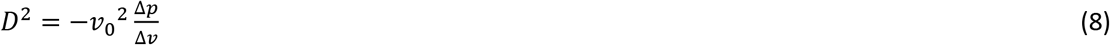

Eq.8 has been referred to as the straight line of Todes(20) as it shows that the initial and final states across a shock jump can be connected by a straight line with the slope 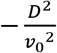 as shown in figure 1. Therefore, initial and final states are strongly constrained by the velocity of the wave front, which also provides the basis for actually measuring the Hugoniots; by ramping up the amplitude slowly and therefore tracing the end of the chord one point at a time. But why do we insist that the Hugoniot has to be different from the adiabatic state diagram for finite amplitudes?

The constraints become apparent when we expand both sides of eq. 5 as a function of *s* and *v*. It can be shown that;

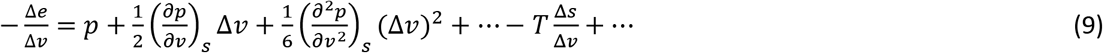

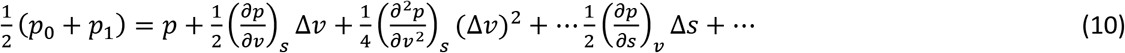

For Δ*v* not infinitesimal but still small 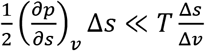 and then comparing eq. 9 and eq.10, which should be equal as per eq.5, we get;

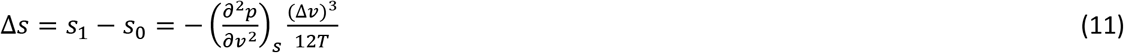

The above equations were derived for the case where a finite amplitude wave moves into material 0 changing into material 1 and this change results from the process that takes place at the wave-front free from any other constraints other than those mentioned above. Therefore, for the wave-front to exist or the process to be possible Δ*s* ≥ 0. The equality is approached for the limit Δ*v* → 0 as discussed above for the infinitesimal acoustic fluctuations. However, for finite case the process will be necessarily irreversible and if the wavefront is compressive then;

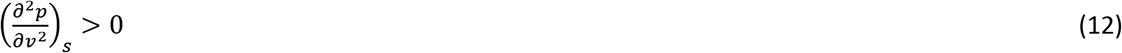

Eq. 11 also quantifies how the Hugoniot diverges from the isentrope with respect to the third power of the relative compression across the wavefront. It can also be appreciated that for same relative compression, the final pressure and temperatures are higher in a shock wave compared to an adiabatic process, because the heat released from the irreversible state changes mostly stays within the shock wave (adiabatic boundary imposed by the speed of sound), which is different from an adiabatic process, where there is no heat transfer even between the subsystems, i.e. no internal heat transfer. Eq.12 is generally fulfilled for all materials as long as only a single phase is present(19), however it breaks near a phase transition as evident from the shoulder that appears in, for example, the (*p*, *v*) phase diagram. Therefore, if the phase change has the time to occur (relaxation time of the phase transition is smaller than the shock compression rate) the wave-front will become unstable. Such considerations acquire fundamental importance in the entire discussion around nerve impulses being a transitional all – or – none wave phenomenon as originally proposed by Konrad Kaufmann. In fact, a significant body of work exists that has explored finite amplitude waves in materials that undergo phase transition (19, 23–26, 30). These past observations have now been replicated for finite amplitude waves in lipid monolayers in detail, which further underlines its importance for the phenomenon of nerve pulse propagation.

## APPLYING SHOCK PHYSICS TO LIPID INTERFACES

Within continuum mechanics, there are two distinct approaches to the problem of stability of wave propagation in an arbitrary fluid; first is based on solving its detailed structure based on Navier – Stokes equations, second is to analyze the relationship between the initial and final thermodynamic state in the inviscid fluid limit across the wavefront, as discussed above. Remarkably, in the second approach, despite the inviscid limit, the viscosities and thermal conductivity effects of a real material are implicitly accounted for in the final results, when both initial and final states are measured experimentally(19). These effects contribute to the smoothening of the wavefront giving it a finite width and rise time, which would otherwise be discontinuous. Therefore, the final state lies on a unique state diagram known as a Hugoniot, measurable only via the wavefront that implicitly accounts for the resulting heat. It was this emphasis on measurement that aligns well with our approach to action potentials and therefore this classical framework was enthusiastically perused in the context of lipid monolayers.

We start with the same principle that predicted the sound like wave-propagation at interfaces in the first place; conservation of the entropy of the interface(31). The entropy of a system is a linear sum of the entropies of its subsystems only where there are no interfaces and no interactions between the subsystems(32). An interface, which is mathematically a two-dimensional system, has an entropy potential(6, 33). Physically the interface has an extension in the third dimension, which in the case of a lipid monolayer consists of the hydration layer and any accompanying ions and solutes. The extent of this dimension is not trivial and depends on both the timescales of a phenomenon(12) and the state, e.g. due to the long-range effects of the surface potential(34). However, instead of focussing on the structure of the medium, we prefer an approach that focuses on the measured state of the interface and accounts for the structure implicitly.

The thermodynamic state of a system in equilibrium can be described as a function of two independent thermodynamic variables. Accordingly, the entropy of the interface can be written as a thermodynamic potential of the form *S*(*E*, *A*), where *E* and *A* are energy and area of the interface. All other variables adjust(including temperature, pH, surface charge and dipole moment, and the width of the interface) such that the entropy is maximum and can be obtained in principle from the complete equation of state. For a system in equilibrium the subsystems are also in equilibrium, which implies that the variables can also represent specific energy *e* and area *a*, i.e. *s*(*e*, *a*) representing per unit mass values. Then by definition, we get an interfacial pressure 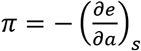, interfacial temperature, 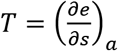 and an interfacial adiabatic compressibility 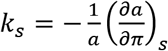. The isothermal state diagrams of the interface consisting of a lipid monolayer have been studied extensively as a classical model for biological membranes(35), in an instrument known as a Langmuir trough, where *π* can directly be measured as function of *A*.

Note that the area *A* controlled by the barriers in an isotherm is different from the area *a* discussed above, which is area per unit mass. The area of a Langmuir trough is sometimes represented in similar units when it us normalised with respect to the amount of lipids added, but we need to be very careful to not use such normalisation here. As discussed, the area appearing in the shock equations represents the reduced mass of the entire 2D region through which the wave propagates, including the water and the ions. As it is not trivial to obtain these number, we will assume that relatively and qualitatively the isothermal state diagram, where area is normalised with respect to the mass of the propagation medium, has the same form as the one observed in a Langmuir trough experiment with some monotonic scaling factor. The form of these state diagrams also depends on changes in temperature, pH, ions, and lipid composition of the system, and anything else that interacts with the interface, including toxins(36, 37), hallucinogens(38), opioids(39), and anesthetics (40). Therefore, even though the control variable is the surface area the isotherm measured on a Langmuir trough doesn’t simply measure a “monolayer” of lipids, it rather measures a quasi-two-dimensional (2D) hydrated region around the interface. The cartoon in fig.2 shows a side view of this interface, where medium I and II are air and water respectively. A representative *π* − *a* isotherm is also shown.

**Figure 2.**
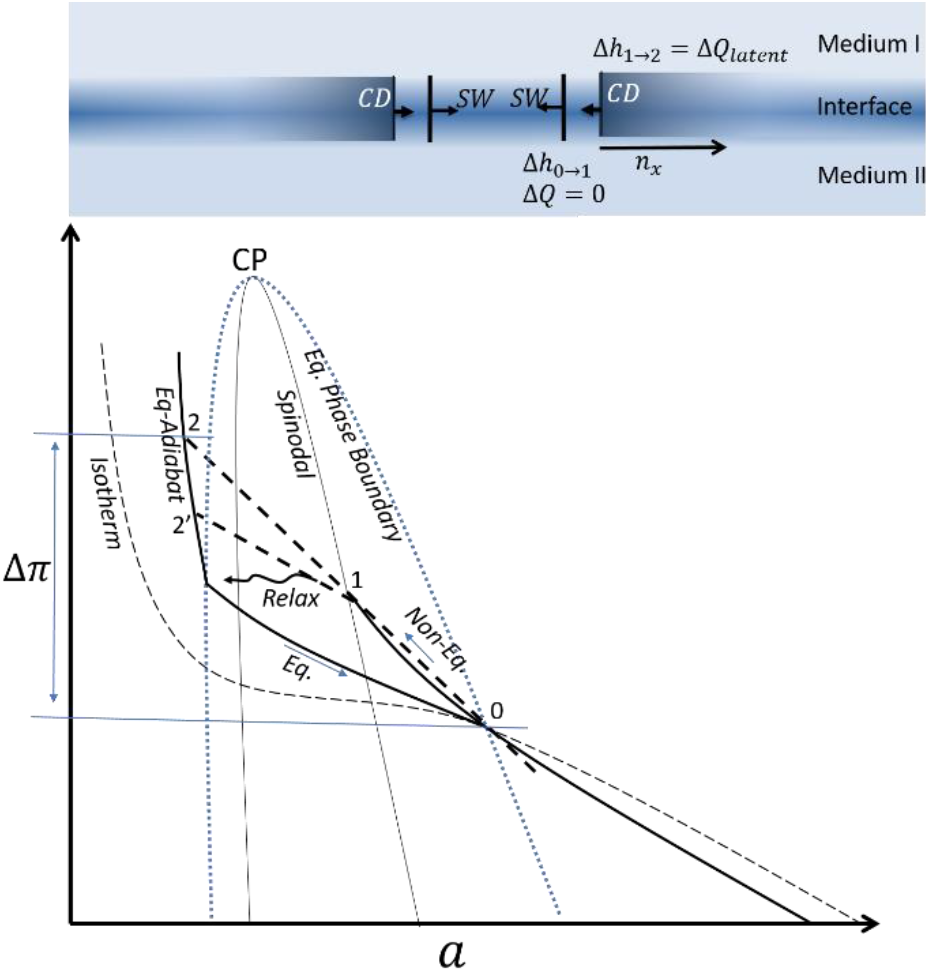
*(Top) Two shock waves followed by their respective contact discontinuities are shown propagating towards each other at an interface. (Bottom) The corresponding jump conditions are shown in the lateral pressure – area or π − a state diagram. The initial state between the two shocks is represented by the subscript 0 and is situated at the isothermal phase boundary. As the condition* 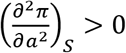 *is violated by the equilibrium isentrope, there is a threshold for excitation and the SW emerges along a non-equilibrium adiabat* 0 → 1, *followed by the phase change* 1 → 2 or 1 → 2’ *across CD. Adapted from* fig.4 in ref. (9))

A local perturbation of the interface away from the state of thermodynamic equilibrium initiates restorative processes including propagating wavefronts. Let the initial state of thermodynamic equilibrium be represented by state (*π*_0_, *a*_0_) on the isotherm. In line with the principle behind Hugoniots, the final state across the wavefront will have to be thermodynamically stable, which is ascertained by its direct observation, and must satisfy the conservation of mass, momentum, and energy, *at the interface.* If we ignore viscosity and thermal conductivity, as discussed above, eliminating flow velocity from conservation of mass and momentum leads to;

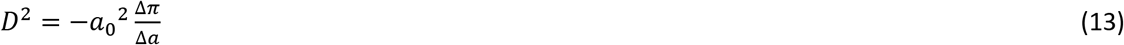

Here, *D* is the velocity of the wave-front 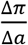, is defined for the relevant thermodynamic process, whereas 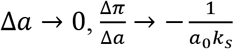, and *D* → *c* i.e. the speed of sound. It should be emphasized that the applicability of adiabatic boundary conditions during sound propagation is not obvious. *Sound waves are adiabatic because the adiabatic properties of the medium give the correct speed of sound, as was shown by the famous correction of Laplace. Therefore, it is crucial to realize that the adiabatic boundary conditions self-organize during sound propagation and the same is true at the interface, even if the physical location of this boundary is not clear.* Indeed *a*_0_ is unknown in absolute terms as it accounts for the effective mass of the self-organized 2D region. However, a detailed solution of the hydrodynamic problem gives its accurate value and correctly predicts the observed wave velocity(12, 41). The equation also indicates that the speed of propagation is directly linked to amplitude, i.e. nonlinearity, and the equation can be plotted as a straight line (dashed lines fig.2) on the *π* − *a* diagram starting at *a*_0_. The other end of the straight line than traces the Hugoniot, while the intermediate points on the straight line have no physical significance.

The second relation is obtained from eq.5 and introducing enthalpy *h* = *e* + *πa*;

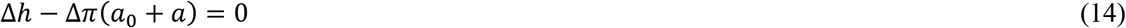

Typically, if the heat changes can be ignored *π* − *a* relation is sufficient to predict several characteristics of these waves correctly, even under nonlinear conditions(12). However, if there is a phase change involved, the latent heat of transition cannot be ignored. The strict condition for an adiabatic process which requires no heat transfer between any subsystem (42) also cannot hold anymore, as the phase change requires a transfer of heat from the lipid tails to the surrounding medium. Accordingly, there is finite relaxation time and as there are no external influences involved the entropy of the system has to increase. In analogy with eq. 11, we can write(19)

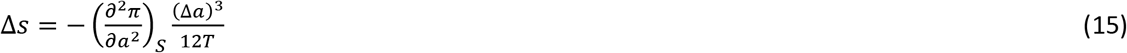

Consequently for a compression shock at an interface to be stable 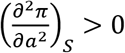. A stable shock wave can be excited either by compressing the medium faster than the relaxation time for phase change, i.e. along with a non-equilibrium path (fig.2) into the superheated regime, or so much that 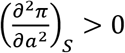 region beyond the phase transition compensates to give an overall Δ*S* > 0. Although it hasn’t be stated explicitly as such before our work to the best of our knowledge, the conditions thus predict a threshold, as indeed previously shown in the lipid monolayer (9). The condition also appears to be related to the observation of the saturation of maximum amplitude of these waves as a function of the excitation strength. The saturation is accompanied by a nonlinear increase in shock width indicating a nonlinear increase in dissipation and dispersion for excitation beyond a limit(14). We believe this is because the medium cannot be compressed arbitrarily deep into the metastable regime. The phase change becomes unavoidable at the so-called spinodal boundary i.e. (*π*_1_, *a*_1_) in fig.2, which limits the maximum slope of eq.13 and hence the maximum amplitude (*π*_2_, *a*_2_). The spinodal decomposition is fundamentally related to the stability of different phase in local equilibrium and occurs when there is no nucleation barrier(43).

## COLLISION OF SHOCK WAVES IN LIPID MONOLAYERS: A CASE STUDY

So, what happens as the super-threshold shock waves slowly dissipate, decreasing the compression rate below the threshold? The intermediate value theorem for Eq.15 between the initial and the final state, implies that the condition 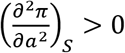 will be violated at some point which should result in a splitting of the shock wave into two waves. The first wave is a shock wave that satisfies eq.15 by avoiding a phase change and compressing the medium into the metastable region given by Δ*π*_0→1_ while Δ*h*_0→1_ ≈ 0. The second wavefront is the so-called contact discontinuity across which phase changes occur and latent heat Δ*h*_1→2’_ is released while Δ*π*_1→2’_ ≈ 0. A contact discontinuity is defined as a surface across which pressure and particle velocity stay the same to first order, but the extensive state variables (entropy, energy, density) change discontinuously, i.e. a phase change takes place (44, 45). The contact discontinuity is thus highly unstable and decays rapidly. However, as the phase change starts at (*π*_1_, *a*_1_) and *c*_1_ > *D*_0→1_ by definition, the released latent heat can propagate partially to the front reinforcing the shock waves maintaining its amplitude and velocity, via a mechanism known as polymorphic detonation(14). It can be claimed that the two wavefronts always exist for a shock wave excited at phase transition, they just co-propagate when the amplitude is sufficiently high given by the slope 0 → 2, and split as the amplitude decreases in two waves 0 → 1 and 1 → 2’. Note that the above discussion is strictly applicable to shock waves, waves excited by a recoiling source as in the lipid monolayer have a more complicated structure (interaction of a shock wave followed by a rarefaction wave^1^). Still, in principle, the entire splitting phenomenon was observed in a lipid monolayer in remarkable detail, which summarises our understanding of the shockwave phenomenon(9) before this article. We have since published the observation of annihilation of two such shock waves upon head-on collision, however without a mechanism, which we now provide.

Let *SW*_→_ be the shock front traveling to the right followed by the contact discontinuity *CD*_→_ traveling in the same direction. Thus as discussed *CD* essentially represents an out of equilibrium phase separation boundary that prefers to be smeared out rapidly compared to the front running shock waves, as observed previously in the lipid monolayer(9). Similarly *SW*_←_ and *CD*_←_ are the two fronts travelling to the left. The cartoon in the inset of figure 2 depicts the situation. Let the initial state, between the shock waves be represented by subscript 0. Then the state across *SW* and *CD* (jump conditions) are represented by the subscript 1 and 2, respectively. The jump condition between these states is shown on the *π* − *a* state diagrams in analogy with the *p* − *v* diagrams in shock physics. While there is an increase in speed of sound across a *SW*, the speed of sound decreases across a *CD* and this relationship is critical in understanding the interaction of these waves.

When a *SW* traveling in a medium of lower sound speed hits a second medium with higher sound speed, it results in a reflected and a transmitted shock wave. Therefore, when *SW*_→_ and *SW*_←_ collide, two shocks waves will immerge from the point of interaction. In general, a new contact discontinuity forms at the collision point, but if the two colliding shock waves are of equal strengths the discontinuity vanishes. Then the transmitted *SW*_→_ meets *CD*_←_, i.e. *SW*_→_ hits a medium with lower sound speed. In this case, two possibilities exist as per the rules of shock wave interactions that depend on the strength of shock wave and the change in the magnitude of the quantity *c*/(1 − *ε*^2^), where 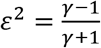 and 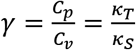 (44). The condition is based on empirical studies that were carried out for a variety of gases in shock tubes and for the theoretical details please see. These empirical studies provided the following criteria: *if c*/(1 − *ε*^2^) *is large and the shock waves are of sufficient strength, a shock wave is reflected and a rarefaction wave is transmitted. However, for a weak shock wave, there will be penetration.*

If we assume that the state of the interface can be estimated locally by a polytropic process, we can apply the above results to collision and the structure of the shock wave discussed above. In lipid systems, we know that the *K*_*T*_ > *K*_*S*_ typically by 10%, however at phase transition *κ*_*T*_ increases dramatically compared to *κ*_*S*_ (46). Therefore the fluid-gel coexistence region that exists behind the *CD*_←_ has a significantly high value of *c*/(1 − *ε*^2^). Accordingly for sufficiently strong *SW*_→_, only a weak rarefaction wave *rW*_→_ is transmitted across *CD*_←_, while the shock wave is reflected. The situation is shown on an (*t*, *x*) plot (fig. 3), typically used to represent such interactions in shocks physics. As shown in fig.3, the shock wave will be trapped by the two contact discontinuities *CD*_→_ and *CD*_←_ resulting in repetitive weakening of *SW*_→_ and *SW*_←_ and weakened shocks would emerge. Following that, the zone between *CD*_→_ and *CD*_←_ will be that of constant pressure and flow velocity, therefore when the discontinuities or the 1D phase boundaries meet they would merge immediately releasing the entropy of the 1D interface, like two bubbles merging together not to be separated, resulting in significant annihilation. The merging is also expected to affect the escaped shock waves *SW* upstream, which were previously seen to be weakened by the disappearance of the downstream *CD* in lipid monolayer experiments (see fig.4 in ref. (9)). In the collision experiments in lipid monolayer(47), it was indeed observed that the degree of annihilation increases with the strength of shock waves, which could be up to 80%. Thus annihilation requires very specific conditions while, in general, interfacial shockwaves would penetrate each other, thus allowing for penetration of neuronal signals, which is also observed in general for excitable systems(48, 49).

**Figure 3.**
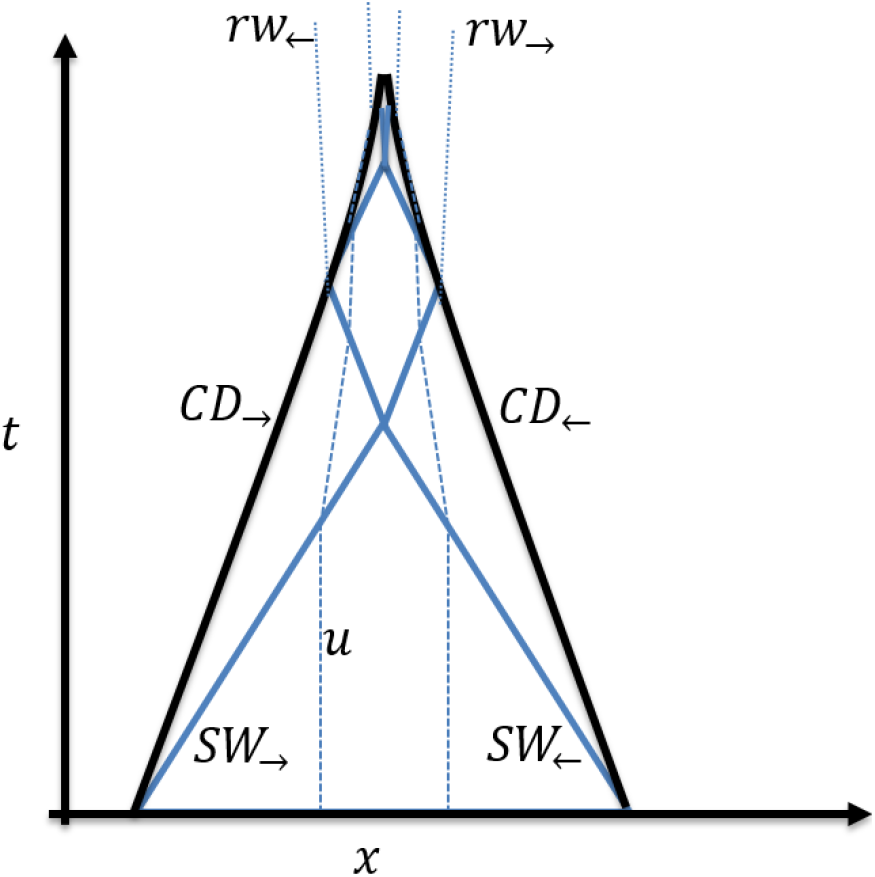
Colliding shocks in space-time. *The figure shows the specific case of annihilation. The path travelled by two shock waves SW_→_ and SW_←_, (blue solid lines) colliding head on followed by their contact discontinuities CD_→_ and CD_←_ (black solid lines). The path of the particles at a given location is also indicated (blue dotted), and the particle velocity u is related to the slope of this path. The slope of the lines changes upon interaction (not all clearly visible), while the slope of the lines CD is non-uniform indicating slowing down and dissipation.*

## SHOCK WAVES VS SOLITONS

Eq. 5 and eq.6 provided the basis for everything we discussed so far and these equations were derived by eliminating the material velocities *u*_0_ and *u*_1_. Consequently, we were not required to solve for the displacement field at any point in the discussion. The solution inherently assumes that all quantities can be represented as function of (*x* ∓ *Dt*) (20), whenever those details are required, which works in 1D. However, as discussed above, it is also possible to solve for the displacement field, for example by solving the Navier-Stokes equation at the interface. The complete solution using this approach requires several inputs that are not directly accessible for a system undergoing phase transition during a wave cycle as all material properties change during the process(50). For example, on one hand the friction between the differential fluid elements, given by the first viscosity, is significantly different between the fluid and the gel phase(51). On the other hand, friction between the degrees of freedom internal to the fluid element (second viscosity(52)), for example due to conformational changes, is also different and significant. Similarly, thermal conductivity and heat capacities also need to be entered, which makes solving the complete problem a formidable task. The shock physics approach overcomes these challenges because it depends on experimentally measured Hugoniots that by definition accounts for these effects implicitly(19). But that also means that Hugoniots based approach can’t be used to construct the solution bottom-up. What the Hugoniot approach instead provides are the macroscopic constraints that a bottom up solution like the one provided by Kappler et.al.(12) or the soliton model by Heimburg and Jackson must satisfy.

When solving the displacement field in terms of (*x*, *t*) it becomes important to think explicitly in terms of space and time. Therefore, the speed of sound is usually represented as *c*(Δ*ρ*, *ω*), indicating that the speed depends both on the compression amplitude (nonlinearity) and frequency (dispersion). However, note that for large compression the higher order terms in the Taylor expansion of *c*(Δ*ρ*, *ω*) are not negligible and the two effects cannot be decomposed as simple nonlinearity and dispersion effects. In Kappler et.al., while the amplitude dependence was estimated from the measured isotherm, the dispersion entered through the first viscosity in the form of coupling between lipids and the surrounding water layer. Therefore, while the model included dissipation effects due to first viscosity, it ignored the relaxation effects (second viscosity) as well as thermal conductivity. The model was able to explain several characteristics of the shock waves in the lipid monolayer quantitatively, including the location of threshold in the phase diagram, i.e. the large amplitude waves were observed only when they crossed the peak in compressibility giving 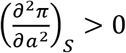, beyond which the velocity increased with amplitude. However, the model was not able to replicate the collision results and threshold and saturation effects were not as strong as those observed in the experiment.

In the soliton model, the origin of nonlinearity was the same, however dispersion was expressed directly in terms of relaxation timescales obtained from ultrasonic exposure of lipid vesicles. The model thus captures the dispersive effects of the relaxation process on the speed of sound but it ignores the dissipative effects of the same phenomenon in the form of second viscosity. Therefore, while in principle it results in a propagating phase transition in line with Kauffman’s original proposition, the details of the solution have some disagreements with the shock theory, which needs further investigation. For example, the soliton model allows completely reversible finite amplitude waves, which as discussed is not allowed in the acoustic theory. Similarly, in a soliton as the amplitude increases the material becomes progressively softer and the speed of sound decreases, i.e. the solution violates 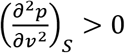. This is in contrast with the waves observed in the lipid monolayers where the waves do not exist till the condition is not met, beyond which the material becomes harder and the speed of sound increases with increasing amplitude. It may be argued that the dispersion relation in the soliton model is not the same as lipid monolayer hence the comparison is invalid and this certainly requires further investigation. However, even in nerves there are evidence of steepening of the wavefront(53, 54), which are consistent with shock physics but not the soliton model. Therefore, while the fundamental premise of the soliton model is consistent with the conservation laws and thermodynamics, the shock physics approach indicates further scope for improvements in the model.

## CONCLUSION AND OUTLOOK

Given that the evidence of a increased order and compression was recently presented for an action potential in a single neuron (55), and an undisputable role of mechanics in neuronal signal transduction(56), the physical insights provided by the above mechanism for collision need to be investigated during the collision or interaction of waves in neurons. In general, excitation waves from propagating graded potentials to action-potential, show an entire range of interactions; from superposition to supra-linear, to sublinear summation(57). Upon interaction, these waves can emerge unaffected(48, 49) to being annihilated(58, 59). Shock physics as developed for lipid monolayer now provides a general mechanism for such interactions at interfaces. In particular, the mechanism of annihilation, in the conventional understanding of action potential and the one based on shock waves, is qualitatively different. Hodgkin and Huxley equations model annihilation on the same basis as the refractory period (60), i.e. how flames propagating from the two ends of a fuse wire stop at the meeting point. However, such a description was categorically refuted by Tasaki’s observations even within the framework of saltatory conduction, quoting, “By collision, transmission of impulses is blocked, not on account of the refractoriness left behind by the impulses, but through lack of internal stimulating current by which the normal transmission is effected”(61). Annihilation of shock waves on the other hand should produce a bang as the momentum and entropy need to be conserved. The phenomenon is therefore analogous to the collapse of a bubble that produces sound (and in extreme cases light(62)). It is however not clear if this emission would escape the nerve or stay within the interface. Based on Tasaki’s measurements of temperature upon collision, it is clear that there is no significant increase in dissipation upon collision, at least none that would be apparent qualitatively(63). A significant increase in dissipation would have resulted in significant distortion of the temperature waveform upon collision, which was not observed. Tasaki did not report quantitatively if the temperature waveform observed during collision was a simple superimposition of individual waveforms without collision, except that they very close. Note that these experiments were performed in nerve fibres and not in single axon and showed significant dispersion. Also it is not clear if the annihilation takes places under all conditions in nerve fibres(64).

Another possibility is the activation of alternate mechanisms that can carry away the momentum and conserve the entropy. For example, an understanding of head-on collision is also important for understanding the fate of shock waves or action potentials as they approach nerve endings, which creates a similar situation as the shock collides with itself (65). Here, in the shock physics framework, the collision can trigger the creation of new interfaces required for the release of neurotransmitters, hence conserving entropy and momentum(66). Clearly, in such cases the boundaries of the system are not contained within the 2D region as discussed so far and it is difficult to track the entropy and momentum. To resolve the difficulty, eq.2 can be used at the terminals along with the approximation 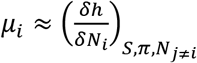, where *μ*_*i*_ is the chemical potential of the expunged *N*_*i*_ molecules of an species *i* from the interface (27), defined conveniently at constant *S* and *π*. Inversely, this line of inquiry will also provide insights into the mechanisms of generation and sustenance(66, 67) of action potentials within the thermofluids and shock physics framework.

What shock physics or thermo-fluids framework allows is a seamless integration of irreversible and reversible aspects of the phenomenon. In fact, we believe that irreversibility plays an important role in the long terms effects of these waves and in enabling adaptation and memory effects as discussed previously(47). The tools provided by shock physics framework might also play an important role in integrating exothermic and endothermic events at the interface, such as opening of channels or action of enzymes, with the propagating steady state shock wave in the membrane. Recent findings on the regenerative role of Nodes of Ranvier provide intriguing possibilities for how to incorporate such phenomenon in an acoustically propagating wave-front, a detonation wave, if such regeneration indeed turns out to be necessary(67). The history of detonation waves in fact provides an excellent case study into how to resolve the acoustic nature of a phenomenon. The question whether the chemical front leads the shock wave or the shock wave leads the chemical front in a detonation wave was a hotly debated topic in 1940s (interestingly the same time as major advances in the conventional theory of action potential). The debate was resolved by following the arguments presented in this study and identifying the two different set of solutions; detonation waves which are supersonic and propagate as compressive front, the other as deflagration waves (flame fronts) that are sub-sonic and propagate as expansion front.

The following statement from Zeldovich was significant in this regard, *“Finally, entirely inadmissible at the present time are the attempts to identify the velocity of detonation with the velocity of motion of any particular molecules, atoms, or radicals in the products of combustion, the corresponding particles being assumed active centers of a chemical reaction chain. However good the numerical agreement, such an attempt is no more than a make-shift and a clear backward step with respect to the thermodynamic theory as is evident from the fact alone that it is entirely unclear what mean or mean square velocity, or other velocity of the molecules, should enter the computation.”*

In contrast, the debate around the acoustic nature of action potentials has predominantly relied on the reversibility of temperature pulse. While a valid argument, it leaves scope for ambiguity as critics have provided irreversible mechanism to explain the observation(2). We can learn from how the acoustic foundation of the detonation phenomenon was historically established, and rephrase the present debate in terms of *efficiency*, or the ratio of irreversible/reversible heat observed during action potentials. This ratio has been consistently observed to be less than 1:10 where as any model that involves ion-channels as the driving mechanism estimates the ratio to be 1:1, i.e. an order of magnitude difference(68, 69). Therefore, a model that explains the temperature pulse based on irreversible mechanism should aim to get the ratio right. Shock physics based explanation provided in this article is consistent with the criteria as in shock waves, the entropy generation depends on the third power of wave amplitude, where as in a RC circuit, the entropy generation would depend on the second power of the wave amplitude, i.e. an order of magnitude difference. Note that while the acoustic nature of action potential certainly needs to be resolved, other excitation phenomenon that spread much slowly, in a reaction diffusion manner, would belong to the deflagration class, and shock physics can provide a universal framework to explain and analyse the two kinds of phenomenon with the same tools.

Finally, coming back to the mechanisms of pulse propagation in a lipid monolayer, all of its main features are now understood, at least in the context of 1D shock physics. However, there are open questions that might pertain to the coupling of these waves to the surrounding media or a consequence of an absolute upper limit to shock compression (70), which we don’t understand yet. For example, when we kept increasing the excitation strength, even after the pulses in the monolayer reached the saturation amplitude, a new wavefront began to emerge ahead of the shock front (Fig. 1(a) in ref (9)). However, within 1D shock physics, no elementary wave can travel ahead of the shock wave. Similarly, during collision experiments, we observed speeding up and increase in the amplitude of the first wave to reach the detector before the collision, in repeated experiments (see supplementary data in ref. (47)). Interestingly, Tasaki also observed a similar effect during the collision experiments, i.e. when two impulses approach one another, they travel from node to node at a rate much greater than in the ordinary transmission. Similarly, a faster time constant and peak amplitude upon collision were also observed in the dog Purkinje system(65). Further research into biophysical changes during the annihilation(59) and penetration(48) of colliding action potential will answer some of these questions.

## Acknowledgment

Funding from EPSRC as part of INSIGHT start-up grant under Rosalind Franklin Institute that supported the research position is acknowledged. Special thanks to Dr. Konrad Kaufman, as his extended lectures motivated this research. I am also thankful to critical discussions with Prof. Matthias F. Schneider and Dr. Christian Fillafer.

## Footnotes

1 Such excitations are referred to as N-waves and are also related to the unpublished observation of a refractory period of these waves that is strikingly similar to action potential. However, we did not publish these results as the refractory period was comparable to the recoil period of the excitation piezo cantilever, which lowers the confidence given the highly non-linear nature of these interactions.

